# Integration of single-cell RNA-seq data into metabolic models to characterize tumour cell populations

**DOI:** 10.1101/256644

**Authors:** Chiara Damiani, Davide Maspero, Marzia Di Filippo, Riccardo Colombo, Dario Pescini, Alex Graudenzi, Hans Victor Westerhoff, Lilia Alberghina, Marco Vanoni, Giancarlo Mauri

## Abstract

**Motivation:** Metabolic reprogramming is a general feature of cancer cells. Regrettably, the comprehensive quantification of metabolites in biological specimens does not promptly translate into knowledge on the utilization of metabolic pathways. Computational models hold the promise to bridge this gap, by estimating fluxes across metabolic pathways. Yet they currently portray the average behavior of intermixed subpopulations, masking their inherent heterogeneity known to hinder cancer diagnosis and treatment. If complemented with the information on single-cell transcriptome, now enabled by RNA sequencing (scRNA-seq), metabolic models of cancer populations are expected to empower the characterization of the mechanisms behind metabolic heterogeneity. To this aim, we propose single-cell Flux Balance Analysis (scFBA) as a computational framework to translate sc-transcriptomes into single-cell fluxomes.

**Results:** We show that the integration of scRNA-seq profiles of cells derived from lung adenocarcinoma and breast cancer patients, into a multi-scale stoichiometric model of cancer population: 1) significantly reduces the space of feasible single-cell fluxomes; 2) allows to identify clusters of cells with different growth rates within the population; 3) points out the possible metabolic interactions among cells via exchange of metabolites.

**Availability:** The scFBA suite of MATLAB functions is available at https://github.com/BIMIB-DISCo/scFBA, as well as the case study datasets.

## 1 Introduction

Cancer is a heterogeneous, multi-factorial and essentially genetic disease, in which various types of mutations alter the functioning and interactions of genes, causing cancer cells to proliferate in an uncontrolled manner. Despite the plethora of cancer-related mutations, a reduced number of recognizable phenotypic hallmarks [17, 18] have been identified. Furthermore, as recently pointed out, neoplastic growth is driven by the disruption of biochemical pathways and networks not only at the single-cell, but also at the population level [38].

Metabolic rearranging, in particular, is a general feature of cancer cells, which reprogram their metabolism to feed their unrestrained proliferation, as it requires high amounts of energy and building-blocks [41]. The design principles underlying the causative role of metabolism in promoting growth as a function of the nutritional constraints are starting to be understood [6, 35] and the idea of targeting the distinctive features of cancer metabolism has received considerable attention [40]. Unfortunately, recent evidences made clear that a single metabolic program cannot be used to globally define an altered tumour metabolism [3], as cancer cells, even within the same tumour, may cope with the above metabolic requirements by engaging different metabolic pathways [3]. Such variability produces different dependencies on exogenous nutrients, and reflects into heterogeneous responses to metabolic inhibitors [39].

Furthermore, in solid tumours, cancer cells are embedded within the tumour microenvironment (TME), a complex network of fibroblasts, myofibroblasts, myoepithelial cells, vascular endothelial cells, cells of the immune system and extracellular matrix. TME also includes chemical gradients of oxygen and nutrients: the complex interaction of all these elements plays a major role in tumour metabolic heterogeneity [19]. The metabolic interplay that occurs among cancer cells and TME - supported by recent evidence on how malignant cells extract high-energy metabolites (e.g., lactate and fatty acids) from adjacent cells [31] - contributes to treatment resistance [37]. Therefore, effective therapeutic strategy should incorporate knowledge of intra-tumour metabolic heterogeneity and cooperation phenomena within cancer cell populations.

Despite major advances in metabolomics techniques [24], single-cell metabolic analyses are lagging behind [43, 15, 44], mainly because of limitations in working with minute amounts of material. Although some improvements have been made, we are still very far from having high-throughput and high-resolution methods. This is a major problem, because flux distributions are currently estimated from metabolite concentrations retrieved from bulk samples, which often contain intermixed and heterogeneous cell subpopulations, thus overlooking possible cooperation and compensation phenomena.

This limitation might partially be overcome by analyzing a population of cells that have a homogeneous metabolism, for example, by using fluorescence-activated cell sorting techniques to isolate population of cells according to specific physical properties. However, when dealing with populations of cells derived from tumours, it is difficult to assess the relative composition of intermingled cancer cells and of stromal elements within the tumour architecture. Therefore, in order to stratify cancer heterogeneous sub-populations, taken, for example, from biopsies, xenografts or organoids, without knowing *a priori* its composition, quantification of fluxes at the single-cell level is needed.

We have previously shown how Flux Balance Analysis [30] of a population of interacting metabolic networks (popFBA) may in line of principle capture interactions between cells [7]. In popFBA an identical copy of the stoichiometry of the metabolic network of the pathways involved in cancer metabolism was considered for each single-cell in the population. We have also shown that constraints mimicking a gradient of nutrient concentrations can be applied. As countless combinations of individual flux distributions, when summed up, can give as a result the population fluxes, the solution to the problem is undetermined. Thus, the challenge is to reduce the number of possible combinations as much as possible, by adding constraints on the single-cell flux distributions.

Contrary to single-cell metabolomics, the field of single-cell transcriptomics is progressing rapidly [32, 35], and current single-cell RNA-seq techniques (scRNA-seq) allow to achieve an unprecedented resolution in the analysis of intercellular heterogeneity. We here propose to exploit scRNA-seq data to set flux constraints of single-cells in metabolic models of cell populations. To this end, we introduce scFBA (single-cell Flux Balance Analysis), a novel computational framework to characterize the metabolism of heterogeneous cancer cell (sub)populations, by translating single-cell transcriptomes into single-cell fluxomes.

As currently no information on scRNA-seq and extracellular fluxes on the very same samples is publicly available, we here applied scFBA to lung adenocarcinoma (LUAD) patient derive xenograft scRNA-seq, collected in [21], while testing different sets of constraints for the extracellular fluxes. To prove the robustness and applicability of our method, we also applied scFBA on further independent breast cancer datasets collected in [4].

## 2 Approach

The existing constraint-based approaches to integrate (bulk) omics data into metabolic models predict the average flux distribution of a population [26, 29, 42], overlooking possible underlying interactions. Single-cell transcriptome could, in principle, be integrated into these models to obtain single-cell specific metabolic models. However, it is clear that their predictions are deeply affected by constraints on extracellular fluxes [26]. Regrettably, there are no experimental protocols that allow to integrate information of nutrient production/consumption rate and transcriptome/proteome of the very same single cell.

Although some attempts to study the cooperation between different metabolic populations have been put forward [5, 23, 20], mostly focused on microbial communities, these methods require indeed *a priori* knowledge about the specific metabolic requirements and objectives of the intermixed populations. Unfortunately, even though metabolic growth well approximates the metabolic function of a population, we cannot assume that each cell within an *in vivo* cancer population proliferates at the same rate, nor that it proliferates at all. The environmental conditions, as well as the composition of the cancer population may change over time. A major example is given by the different proliferation rates of stem and differentiated cells [1]. For this reason, differently from, e.g., [20], our approach does not assume that the population composition is at equilibrum.

scFBA aims at portraying a snapshot of the single-cell (steady-state) metabolic phenotypes within an (evolving) cell population at a given moment, and at identifying metabolic subpopulations, without *a priori* knowledge, by relying on unsupervised integration of scRNA-seq data. Our idea to achieve this goal is to solve a unique mass balance problem to identify the possible combination of single-cell steady states that concurrently satisfies constraints on single-cell transcriptomes and bulk fluxes.

An identical copy of the stoichiometry of the metabolic network of the pathways involved in cancer metabolism is first considered for each single-cell in the bulk. To set constraints on the fluxes of the individual networks, represented by the single-cell compartments of the multi-scale model, scFBA then distributes the total (population) flux of each reaction proportionally to a score that quantifies the activity of that reaction in each cell.

Similarly to what has been done in [22] for bulk data, the activity scores are computed as function of the expression of the genes encoding for the subunits and for the isoforms of the enzyme catalyzing each reaction in the network. The assumption is that the expression of subunits limits the activity of the enzyme, whereas the expression of isoforms contributes to enhance the activity of the enzyme. We do not use the mRNA levels as a direct proxy of protein levels, because of the many factors that contribute to determine the expression level of a protein [25]. Once the activity scores are computed, scFBA sets the flux upper limit of each reaction in each cell as a fraction of the total possible flux proportional to its activity score.

Because cells within the tumour share the same environment, in order to determine the flux distribution of each network, each cell is not considered individually, but it may interact with any other cell in the population, via release/uptake of metabolites into/from the TME.

A scheme of the scFBA approach is depicted in Figure 1.

**Figure 1:**
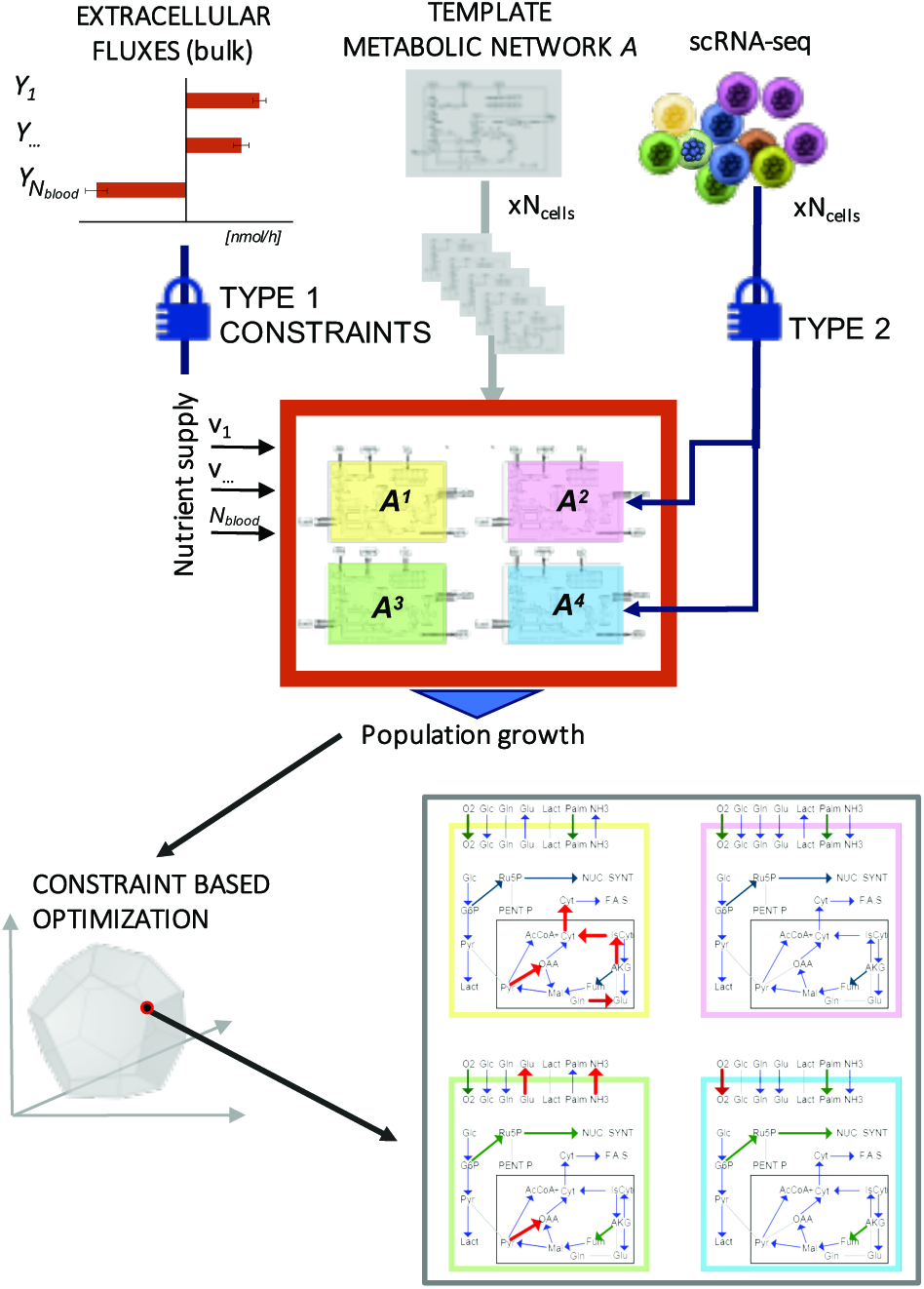
Graphical representation of the scFBA methodology. Extracellular fluxes and single-cell transcriptomes are translated respectively into type 1 and type 2 constraints (see Materials and Methods) of a population of *N*_cells_ replicates of metabolic network *𝒜*. The output is the prediction of single-cell metabolic fluxes.

## 3 Methods

### 3.1 popFBA

Here we briefly recall the popFBA approach. For a more comprehensive description of FBA, the reader is referred to [7].

Starting from a template metabolic network model A, corresponding to a generic single-cell and defined as *A* = (*χ*^*A*^, *ℛ*^*A*^, *ε*^*A*^) - where *χ*^*A*^ = {*X*_1_, …, *X*_*M*_} is the set of metabolites in network *A*, *ℛ*^*A*^ = {*R*_1_, …, *R*_*N*_} the set of biochemical reactions taking place among them and *ε*^*A*^ = {*E*_1_, …, *E*_*N*_*ext*__} is a set of *N*_ext_ unbalanced reactions (exchange reactions), enabling a predefined set of metabolites (including the pseudo-metabolite representing biomass) 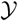 = {*Y*_1_, …, *Y*_*N*_ext__} ⊂ *χ*^*A*^ to be inserted in or removed from the system - the popFBA procedure first builds a population model composed of *N*_cells_ replicates *A*^*c*^ of network *A*, each one corresponding to a single-cell *c*, *c* = 1, …, *N*_cells_, and which can cooperate by exchanging nutrients in the tumour microenvironment.

For each single-cell *c*:

- *A*^*c*^ = (*χ*^*c*^, *ℛ*^*c*^, *C*^*c*^) is its metabolic network;
- *χ*^*c*^ = *χ*^*A*^ is the set of its metabolites;
- *ℛ*^*c*^ = *ℛ*^*A*^ is the set of its internal reactions;
- *𝒞*^*c*^ = {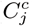}, with *j* = 1, …, *N*_ext_, is a set of cooperation reactions, defined as reactions that allow to exchange metabolites among single-cells via a shared environment that represents the TME compartment. Cooperation reactions are built by transforming each exchange reaction *E*_*j*_ *∊ ε*^*A*^ into a cooperation reaction 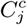 with the form:

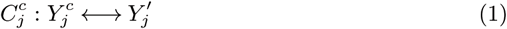

Accordingly, a new set of metabolites pertaining to the TME compartment 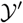 = {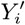} with *i* = 1, …, *N*_ext_ must also be defined.

Because original exchange reactions have been replaced by cooperation reactions, a new set of *N*_blood_ exchange reactions 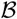 = {*B*_1_, …, *B*_*N*_blood__} is defined, which allows a subset of metabolites 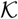 = {*K*_1_, …, *K*_*N*_blood__} ⊂ 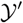 to be exchanged with the external environment, e.g., the blood supply:

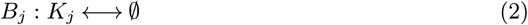

The population model *P* is then defined by: (*i*) the union set of the metabolites *χ*^*P*^ = ⋃_*c*_ *χ*^*c*^ ∪ 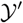; (*ii*) the internal reactions *ℛ*^*P*^ = ⋃_*c*_ *ℛ*^*c*^; (*iii*) the cooperation reactions *𝒞*^*P*^ = ⋃_*c*_ *𝒞*^*c*^; (*iv*) the population exchange reactions 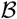.

A stochiometric matrix *S*^*P*^ is then built for all reactions in *ℛ*^*P*^, *𝒞*^*P*^ and 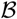 and for all metabolites in *χ*_*P*_ and 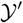. The final size of matrix *S*^*P*^ is (*N*_cells_ · *M* + *N*_blood_) x (*N*_cells_ · (*N* + *N*_ext_) + *N*_blood_). Once the population model is obtained, the total biomass of the *N*_cells_ single-cells is maximised by means of linear programming, as in standard FBA [30].

The solution of popFBA represents the flux distribution

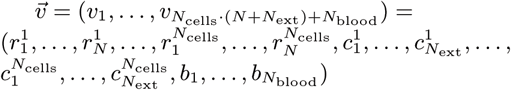

that maximises the biomass exchange flux *b*_*biomass*_, with *v*_*i*_ representing any flux *i* of the population model, and for each single-cell *c*, 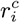 representing the *i*-th internal flux, 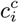 representing the *i*-th cooperation flux and *b*_*i*_ an exchange flux with blood. The optimization problem is postulated as follows:

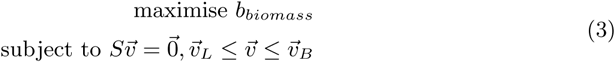

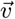_*L*_ and 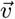_*B*_ are vectors specifying the lower and upper bound respectively for each flux *v*_*i*_ of 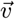. A negative lower bound indicates that flux is allowed in the backward reaction. To solve the above problem we exploited the Gurobi solver within the COBRA Toolbox [33].

### 3.2 Input and data pre-processing

scFBA takes as input a scRNA-seq dataset in the form of a *N*_genes_ x *N*_cells_ matrix *T*, where *N*_genes_ is the number of genes and *N*_cells_ is the number of single-cells under study. Each element *T*_*g*,*c*_, *g* = 1, …, *N*_genes_, *c* = 1, …, *N*_cells_ corresponds to the normalized read count of gene *g* in cell *c* such as, for instance, the TPM (Transcripts Per Kilobase Million).

In addition to scRNA-seq data, certain datasets include RNA-seq profiles of control bulk samples (see, e.g., the datasets described in Section 3.5). scFBA allows to employ the information on bulk expression profiles to make the analysis more robust against possible data-specific errors of single-cell datasets. In particular, scFBA pre-process the data in order to identify genes that might show inconsistencies between bulk RNA-seq and scRNA-seq profiles, by focusing especially on genes with a null read count: for genes displaying a RNA read count equal to zero in all single-cells, but greater than zero in the bulk, we replace the read count for that gene in each cell with the bulk read count. As it will be mentioned below, this step will prevent the erroneous removal of relevant reactions.

### 3.3 Reaction Activity Scores

In order to convert the information on single-cell transcriptomes into constraints on the fluxes of a popFBA model, we assume that enzyme isoforms contribute additively to the overall activity of a given reaction, whereas enzyme subunits limit its activity, by requiring all the components to be present for the reaction to occur.

To this end, we define a Reaction Activity Score (RAS), for each single-cell *c* = 1, …, *N*_cells_, and each reaction *j* ∊ *R*, based on Gene-Protein Rules (GPRs). GPRs are logical formulas that describe how gene products concur to catalyze a given reaction. Such formulas include AND and OR logical operators. AND rules are employed when distinct genes encode different subunits of the same enzyme, i.e., all the subunits are necessary for the reaction to occur. OR rules describe the scenario in which distinct genes encode isoforms of the same enzyme, i.e., either isoform is sufficient to catalyze the reaction.

In order to compute the RAS we distinguish:

- Reactions with AND operator (i.e., enzyme subunits).

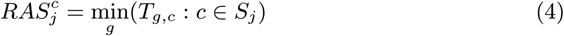

where *S*_*j*_ is the set of genes that encode the subunits of the enzyme catalyzing reaction *j*.
- Reactions with OR operator (i.e., enzyme isoforms).

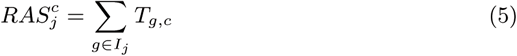

where *I*_*j*_ is the set of genes that encode isoforms of the enzyme that catalyzes reaction *j*.

In case of composite reactions, we respect the standard precedence of the two operators.

### 3.4 scFBA

The first step of the scFBA approach is the creation of a population-specific model of the metabolic network *A*, by removing from the set of reactions *ℛ*^*A*^ those reactions that are expected not to occur in any cell within the population under study, because the competent enzyme is not expressed.

To this end, after processing the expression dataset as described in Section 3.2, we identify the set *G*_*off*_ of genes that, with a high degree of certainty, are not expressed in any single cell (read count after processing equal to 0 in each cell). We then perform a canonical gene deletion of all genes in *G*_*off*_ (Cobra Toolbox [33] function: *geneDeletionAnalysis*), which results in removal of reactions for which their expression is essential.

Once the population-specific model is obtained, the scFBA approach imposes two kind of constraints:

> *type 1* constraints on the extracellular fluxes of the overall population model *P*, i.e., the upper and lower bound of the *N*_blood_ exchange reactions in set 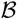, ideally according to metabolic measurements;

> *type 2* constraints on internal fluxes of each single-cell *c*, i.e. for the reactions in *ℛ*^*c*^, with *c* = 1, …, *N*_cells_, and for its set of cooperation reactions *C*^*c*^, according to their RAS, whenever the computation of a RAS is possible, i.e., when a GPR exists for the reaction, along with the transcript values of at least one of the involved genes.

In order to translate the information of the activity score of a given reaction *j*, in a given cell *c*, 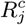:

- we first estimate the possible flux that reaction 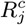 can carry, when only constraints on extracellular fluxes (type 1) are set, whereas the internal fluxes (type 2) are still unbound and the system is not required to make biomass, i.e., we compute the maximal flux in both the forward (*F*_*ƒ*_) and backward direction (*F*_*b*_) of each reaction. To do so, we perform a Flux Variability Analysis [27], with no optimality required (Cobra Toolbox [33] function: *fluxVariability*). We define 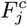 = max(|*F*_*ƒ*_|, |*F*_*b*_|).
- we then compute the relative reaction activity score of 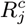 in each *c* = 1, …, *N*_cells_, with respect to the total activity of reaction *j*, as follows:

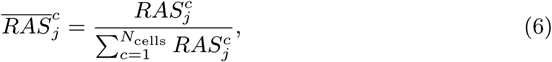
- finally, we assign an upper bound (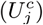) to reaction 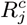, as portion of 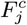 which is proportional to the activity score (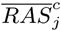) of reaction 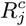. Namely, we remap the values 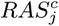, *j* = 1, …, *N*;*c*=1, …, *N*_cells_ in the interval [*∊*, 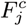], as follows:

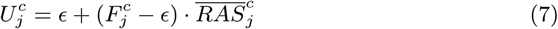

We set the upper bound of reactions having 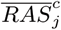 = 0 to a small value *∊* rather than to 0 to mitigate the impact of false-negatives. We remark that genes that have no read count also in the bulk sequencing are instead fully deleted from the model. In this study we set *∊* = 10^−3^. Note that 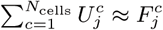.
- if reaction 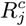 is irreversible, we assign a zero lower bound (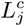 = 0) to reaction 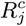, otherwise we assign a lower bound 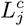 = −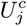.

The reason why, when dealing with reversible reactions, we avoid setting different values for backward and forward reaction, by assigning to 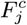 the maximum value between *F*_*ƒ*_ and *F*_*b*_, is that we should be agnostic about the preferred direction, given that the RAS reflects the gene expression of its competent enzyme, which may work in either direction.

Once the *P* model is constrained (with both type 1 and type 2 constraints), Linear Programming, as well as other standard constraint-based methods can be applied.

### 3.5 Datasets

In this work, we mainly use the following 3 LUAD datasets obtained from the NCBI Gene Expression Omnibus (GEO) data repository under accession number GSE69405.

LCPT45 Composed of 34 cells acquired from a surgical resection of a 37-mm irregular primary lung lesion in the right middle lobe of a 60-year-old untreated male patient. A xenograft was then obtained by a sub-renal implantation in mice.

H358 Composed of 50 cells from NCI-H358 bronchioalveolar carcinoma cell line.

LCMBT15 Composed of 49 cells acquired from a surgical resection of a metachronous brain metastasis acquired from a 57-year-old female after standard chemotherapy and erlotinib treatments. A xenograft was then obtained by a sub-renal implantation in mice.

We repeated all the analyses also on the following two independent breast cancer datasets (GEO access number: GSE75688), including scRNA-seq data of single-cell suspensions of cancer tissues obtained on the day of the surgery of untreated breast cancer patients [4]:

> BC04 Composed of 59 human epidermal growth factor receptor 2 positive (HER2+) cells.

> BC03LN Composed of 55 lymph node metastases of human estrogen receptor positive (ER+) and (HER2+) cells.

Each of the 5 datasets includes the gene expression level of more than 2000 genes in the form of Transcript Per Kilobase Milion (TPM). We filtered out a few cells with less than 5000 genes detected. For each dataset, we retained only the metabolic genes included in HMRcore model (418 genes). The dataset transcripts are identified by Ensembl ID, which we automatically converted into HUGO Gene Nomenclature Committee (HGNC) ID. Datasets also contain the expression profile of the bulk samples, which we used to pre-process data as described in Section 3.2.

### 3.6 Metabolic network model

In this study we used, as template network A, the metabolic core model HMRcore introduced in [12] and used in [7]. As exchange of fatty acids between cells in tumours has been recently reported [2], we included the possibility to exchange palmitate via the TME and, accordingly, mitochondrial palmitate degradation and gluconeogenesis. Given the importance of reactive oxygen species (ROS) metabolism observed in [6], we also inserted ROS production and removal pathways. As the original version of the model does not include information on GPRs, such rules have been extracted from Recon 2.2 [36] and included in the HMRcore model, and recently manually curated [16]). We decided to disregard the GPR associated to the complexes I to IV of the electron transport chain in scFBA computations, because it unrealistically requires up to 81 genes (AND rule) and we were not able to accurately tune the rule for these elaborate complexes in [16]. However the flux trough complexes I to IV should be modulated by the constraints on the last step of the chain (ATP synthase).

The final version of the HMRcore model includes 315 reactions (of which 263 are associated with a GPR) and 418 metabolic genes. The SBML of the model is provided in https://github.com/BIMIB-DISCo/scFBA.

**Experimental setting** As baseline experimental setting, we considered as main exogenous nutrients (which the overall population can uptake from 0 up to Npop · 100 nmol/h) those used in [7]: glucose, glutamine, oxygen and arginine. As in [7], we considered a cooperation reactions for: glutamate, NH_3_ and lactate; plus palmitate. As a first approximation, we defined as set of nutrients that can be secreted by the population the same set of nutrients considered for the cooperation reactions.

## 4 Results

### 4.1 Integration of RNA-seq data efficiently reduces the space of optimal solutions

We first applied scFBA to the 5 datasets described in Section 3.5, assuming maximization of total (population) biomass production rate as objective function. Worth of note, all five population models display a non negligible maximal growth rate, something that cannot be taken for granted when integrating transcriptomics into FBA models [26, 29, 42].

As a first result, we illustrated that, by integrating scRNA-seq data into the standard population model, the scFBA approach efficiently reduces the space of optimal solutions. To this end, we compared the variability of the biomass production flux of each of the *N*_cells_ single-cells simulated within the population model, for each of the 5 datasets under study.

When no information on cells’ transcriptome is employed (as in standard popFBA settings [7]), the type 2 constraints of the metabolic network are identical for each cells. This implies that each cell is capable of contributing alone to 100% of the objective function value (i.e., the biomass of the total population). As depicted in Figure 2A-E (left plots), the biomass flux value of each cell, within the set of optimal solutions, spans indeed from 0 to 100% of the total biomass (purple rectangles). On the contrary, after scRNA-seq data integration, as performed via scFBA, the biomass flux of each cell can take a specific value, corresponding to a certain fraction of the total biomass (green rectangles, which results in a single line, because the maximum and minimum flux values coincide.)

**Figure 2:**
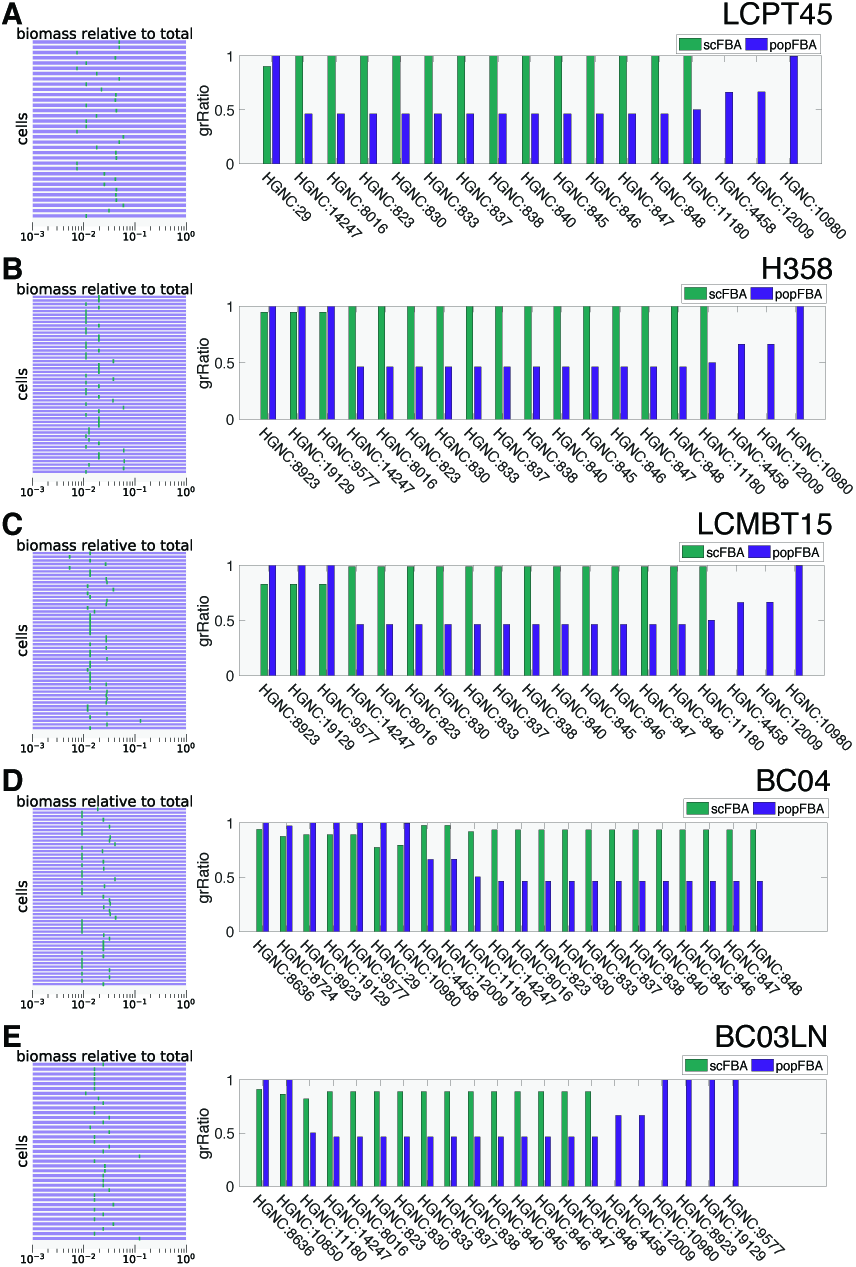
scFBA vs. popFBA. A) Dataset LCPT45. Variability of the fraction of the biomass synthesis flux (logarithmic scale) for each cell over the population growth rate (left panel) before (purple) and after data integration (green). Effect of gene deletion (bars in right panel) on the population growth rate before and after data integration. When *gr Ratio* = 0 (lethal gene), the corresponding bar is not displayed. B-E) Same information as in A for different datasets. In order: H358, LCMBT15, BC04 and BC03LN.

To show how this reduction in the space of alternative optima may actually affect predictions, we performed a single gene deletion analysis with and without scRNA-seq data integration (scFBA and popFBA, respectively). When a single gene is deleted, the reactions for which the expression of such gene is essential (i.e., reactions exclusively associated to the gene, or reactions associated to that gene and other genes with an AND operator) are removed from the network (i.e., from the set *ℛ*^*c*^, ∀c). After removal, the population model is newly optimized for total biomass production, and the growth ratio (*grRatio*) of the new biomass over the previous one is computed. Figure 2A-E (right bar plots) reports the *grRatio* observed for those genes deletions that display a different effect before and after data integration. Notice that when the *grRatio* equals 0, the corresponding bar is not displayed at all.

Remarkably, some genes that are irrelevant (*grRatio* = 1) in popFBA settings (i.e., with no data integration) may become even lethal in scFBA settings (i.e., with scRNA-seq data integration) (*grRatio* = 0). This is the case, for instance, of gene HGNC:10980, which encodes for enzymes responsible for glutathione/phosphate, fumarate/phosphate or α-ketoglutarate/malate antiports. Conversely, some genes that display a significant effect (*grRatio* ≈ 0.5) in popFBA become instead irrelevant in scFBA. This is the case of the genes that encode for ATP synthase (HGNC:823, 830, 833, 837, 838, 840, 845-848, 14247, 8016), indicating that the integration of scRNA-seq data forces a flux distribution in which such reactions carry a suboptimal flux, thus resulting in a milder effect when the reaction is depleted.

### 4.2 scFBA extracts useful features from transcript signals

Single-cell fluxes are expected to be less noisy than transcript signals, which are typically analyzed by means of multi-variate statistical analysis [35] and, therefore, might be used to better identify cell clusters that might represent distinct metabolic subpopulations. To investigate this issue, we performed a cluster analysis on the expression values (scRNA-seq) of the metabolic genes and compared the results with those of a cluster analysis performed on the fluxes predicted by scFBA. To this end, we performed both hierarchical and k-means cluster analysis. In order to avoid reactions with typical high flux-value to induce a bias on clustering results, we first remapped the flux values of each reaction *j* in the interval [0,1]: value 0 is assigned to the cell showing the lowest value for a given flux, 1 to the the one showing the highest value.

Figure 3 reports, for each dataset under study (panels A-E), the results of the hierarchical clustering analysis (distance metric: euclidean), for transcripts (left column) and fluxes (middle column). One can see, from the dendrograms and heat maps, that cells cluster in a few well-separated groups, when the extracted features (the fluxes) are considered, whereas they cluster in many “singletons”, when the original features (the transcripts) are used. For example, when observing the fluxes computed for the dataset LCPT45 (panel A), it is apparent that two major groups of cells can be identified, corresponding respectively to the blue and red-coloured leaves in the dendrogram. We verified that these two groups significantly differ in their growth rates (Z-scores: 3.23; P-value: 0).

**Figure 3:**
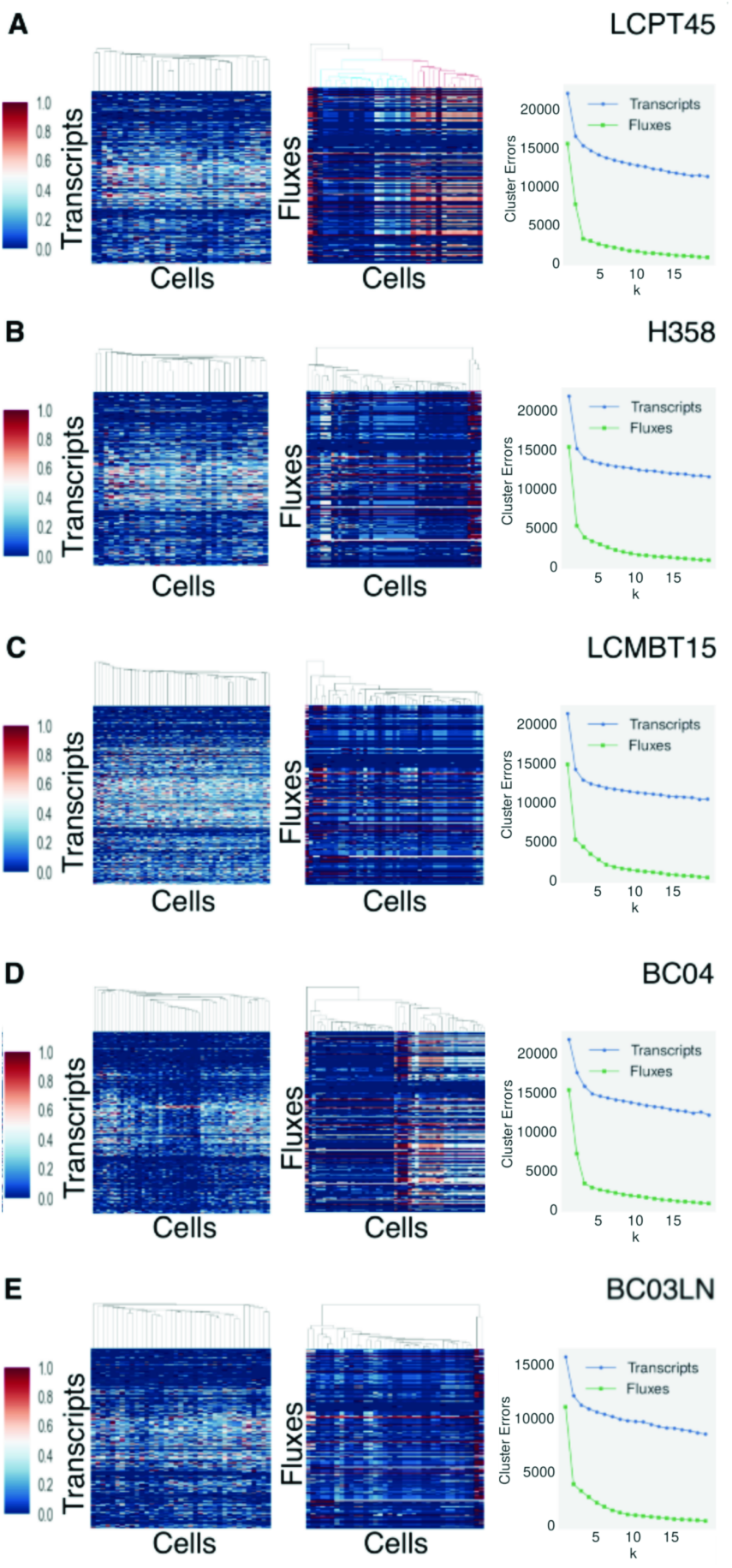
Clustering of transcripts vs. fluxes and elbow analysis. A) LCPT45 dataset. Cluster-gram (distance metric: euclidean) of the transcripts of the metabolic genes included in metabolic network (left panel) and of the metabolic fluxes predicted by scFBA (middle panel). Right panel: elbow analysis comparing cluster errors for *k* ∊ {1, …, 20} (k-means clustering) in both transcripts (blue) and fluxes (green). B-E) Same information as in A for different datasets. In order: H358, LCMBT15, BC04, BC03LN.

To quantitatively compare the clustering of transcripts and fluxes, we first performed a k-means clustering with different number of clusters *k*, by considering *n* = 100 bootstrap iterations (with random centroid assignments) and by selecting the clustering resulting in the maximum inter-cluster distance. We then assessed the clustering goodness, by means of “elbow” and “silhouette” evaluation methods.

The elbow method looks at the variance explained as a function of the number of clusters. It assumes that the number of clusters to be identified should be the smallest *k* able to provide a good modeling of the data (i.e., adding a further cluster does not improve the minimization of the sum of squared errors, SSE). When plotting *k* versus the SSE (Figure 3A-E, right column), for small values of *k* there is typically a rapid decrease of the SSE; however at a certain point the marginal gain in explained variance drastically decreases (an elbow in the plot). The correct number of clusters is identified at the elbow. For instance, in the right column of Figure 3, it is possible to notice that, for the fluxes relative to the primary tumour datasets (panels A and D), a sharper elbow is observed at *k* = 3, hence the optimal number of clusters is 3. Moreover, when comparing transcripts and fluxes datasets, it can be observed that the “cluster errors”(i.e. SSE) are always higher for the former.

The silhouette score estimates for each element the cohesion (similarity of an object with
other elements of its own cluster) and the separation indexes (dissimilarity with elements of other clusters). The silhouette has values in [−1,1]: scores close to 1 indicate that an element is correctly clustered, whereas scores close to −1 indicate a wrong cluster assignment. A silhouette value around 0 indicates that the element is located among two different clusters. In Figure 4, we evaluated the silhouette for the dataset LCPT45 transcripts (A) and fluxes (C) for *k* = 3, i.e., the value identified from the “elbow” analysis, which also corresponds to the highest average silhouette value, when varying *k* in {2, …,6} (data not shown). Noteworthy, in the fluxes case the average silhouette value is considerably higher than the transcripts case (where the score is almost 0) indicating that the calculation of fluxes leads to a better clustering as compared to the evaluation of transcripts. In order to highlight the better subdivision in clusters of dataset fluxes, in Figure 4 we also plotted the feature space for the first two features. From the plot it is possible to notice that in dataset fluxes (D), cluster centroids (circled numbers), have a better separation as compared to transcripts (B).

**Figure 4:**
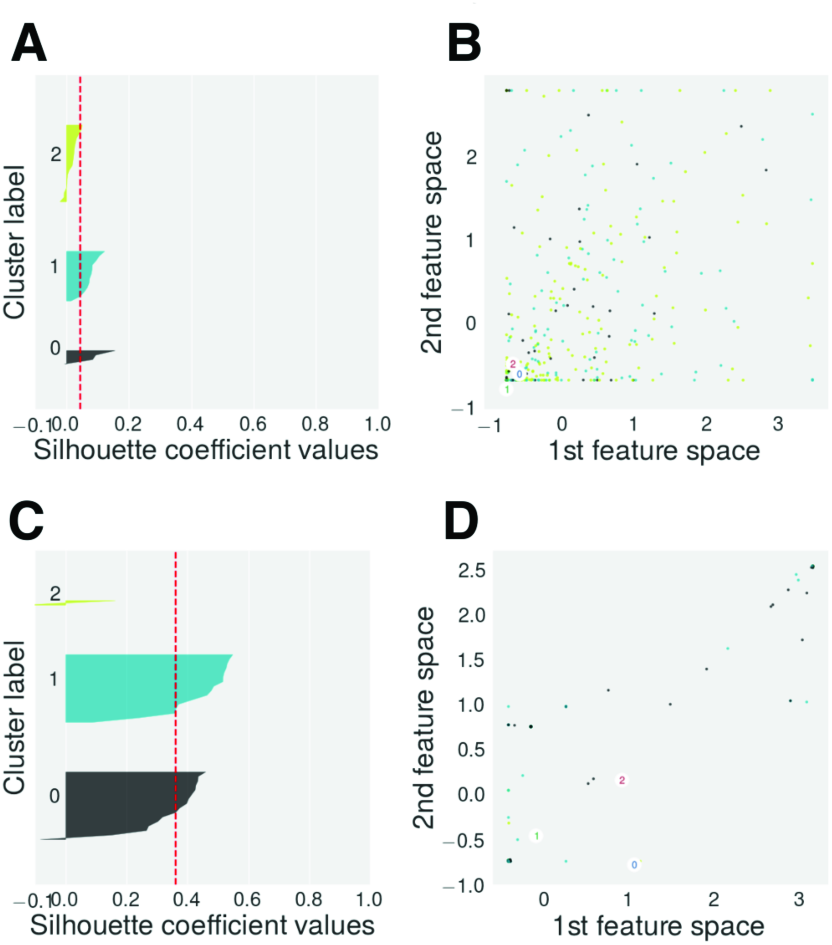
K-means cluster validation of transcripts vs. fluxes. A) Silhouette analysis for LCPT45 transcripts, when *k* = 3. Red dashed line indicates the average silhouette for the entire dataset. B) 1st and 2nd feature space plot for LCPT45, circled values indicate centroids for the *k* = 3 clusters. C-D) Same information as in A and B for LCPT45 fluxes.

All in all, the results of the cluster analyses indicate that fluxes can be better clustered than their transcript counterparts. Remarkably, it can also be noticed (see the sharper “elbow” and the clustergram in Fig. 3A) that the primary tumour xenograft (LCPT45) fluxes better partition into clusters as compared to the cell line (Fig. 3B, H358) and to the secondary tumour xenograft (Fig. 3C, LCMBT15). This result is in line with the data reported in [21], indicating that the presence of heterogeneous populations is emphasized in LCPT45. Notably, we detected a similar difference between the clustering results of primary and secondary tumour of the independent breast cancer datasets (BC04 and BC03LN). Indeed, the former

### 4.3 scFBA captures interactions between cells

The main rationale behind solving a unique mass balance problem for many cells together, given constraints on the extracellular fluxes of the bulk, rather than many separate mass balance problems, is that the nutrient consumption and secretion rates (extracellular fluxes) can be easily determined or approximated from measurements of the concentration of metabolites in the cell culture media at different time points for the bulk only. Another major side benefit of this approach is that it allows to identify the possible interactions among cells within a population, as pointed out in [7].

We verified that, after data integration, some cells secret metabolites that are up-taken by other cells. The heat map in Figure 5 shows the (normalized) flux values of cooperation reactions for the LCPT45 dataset: a positive value means that the cell is secreting the metabolite in the tumour microenvironment, whereas a negative flux that the cell is up-taking it from the tumour microenvironment. It can be observed that a complex network of interactions is established among cells. In particular, a consistent group of cells consumes the lactate and palmitate that are secreted by other groups. The scatter plots in Figure 5 show the dispersion of the fluxes of uptake/secretion from/into the TME for lactate and palmitate and how they couple with different growth rates, portraying a relationship far more complex than that depicted with popFBA (no scRNA-seq integration [7] and no exchange of palmitate allowed).

**Figure 5:**
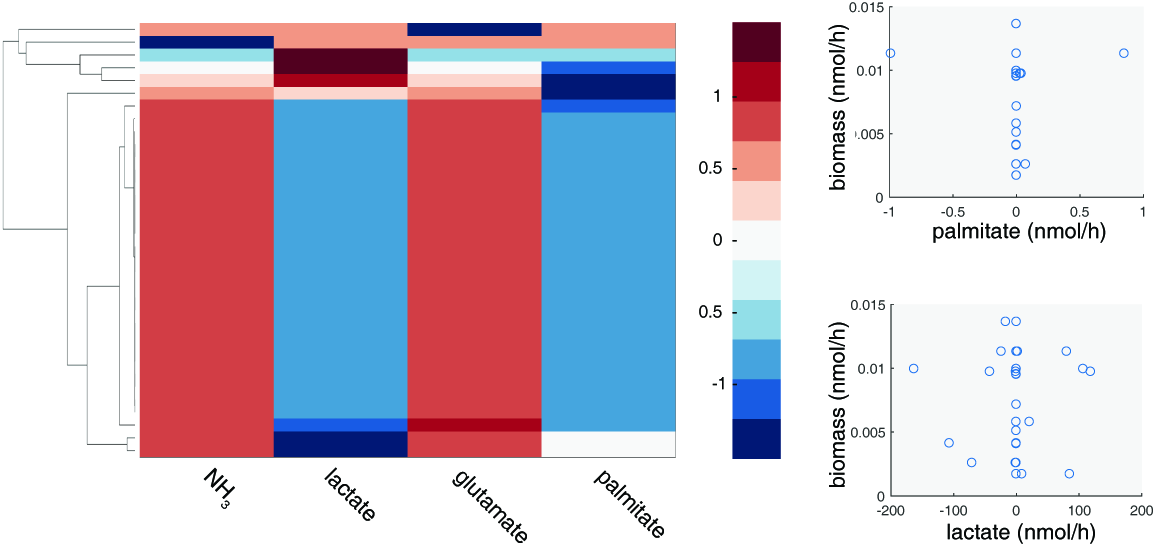
Metabolic cooperation in LCPT45 population. A) Clustergram of the fluxes of cooperation reactions for NH_3_, lactate, glutamate and palmitate. Negative fluxes (blue shades) indicate an uptake, whereas positive fluxes (red shades) indicate a secretion of the corresponding metabolite. B) Scatterplot of the biomass flux values of each cell in the population vs. palmitate (left panel) or vs. lactate cooperation flux (right panel).

#### Effect of cooperation on growth

The optimal values for the cooperation fluxes reported in Figure 5 display a larger variability than the optimal growth rates (Fig. 2), even though much lower as compared to popFBA settings (by at least 60%). Therefore, we verified that the interaction among cells, given their transcriptomes, improves the capability of the overall population to achieve metabolic growth, while also correcting for the possible presence of thermodynamically infeasible loops [11]. At this aim, we compared the population growth rate of the case in which cooperation reactions are allowed, with the case in which they are not, given the same constraints (type 1 and type 2). As the mere secretion of metabolites (such as lactate) in the external environment (e.g., the blood) can improve growth rate, under given boundary conditions (e.g., limiting oxygen [6]), in order to allow for a meaningful comparison, in our experimental setting metabolites that can be secreted in the TME can also be secreted directly in the blood supply. By doing so, when the cooperation reaction is removed, the cell can still rid off of exceeding metabolites, which cannot however be taken up by other cells.

Remarkably, we observed that the ratio of the total biomass obtained in the absence of cooperation reactions over that in their presence may be lower than 1, implying that removal of cooperation limits the capability to achieve growth. In particular, we observed a ratio of: 0.90 for the LCPT45 dataset; 0.99 (H358); 0.99 (LCMBT15); 0.76 (BC04); 0.95 (BC03LN).

Intriguingly, but not surprisingly, the impact of cooperation prevention is higher on those datasets corresponding to more heterogeneous populations (LCPT45 and BC04). Intuitively, cells specialized in different metabolic programs are more likely to cooperate, as compared to similar cells.

#### Effect of cooperation on ATP production

For the sake of simplicity, in this study we assumed an optimal growth rate for the overall population, yet other assumptions may be easily investigated with the scFBA approach. Among others, it is common practice in constraint-based modeling to optimize for ATP production [13, 34]. As a proof of principle, we repeated the analysis on the effect of cooperations when the objective function is the total ATP produced by the population. We obtained the following ratios for the 5 datasets: 0.77 (LCPT45); 0.97 (H358); 0.93 (LCMBT15); 0.99 (BC04); 0.87 (BC03RLN). The observed discrepancies in the extent of the effects of cooperation inhibition on growth and energy productions are worth of interest and would deserve further investigation. Notably, both the energy production and growth rates of the H358 (cell line) population, which is expected to be homogeneous, are not affected by cooperation prevention.

### 4.3 Boundary conditions affect scFBA predictions

We have shown how the integration of scRNA-seq data greatly reduces the space of feasible FBA solutions, meaning that scRNA-seq encodes information on how nutrient utilization should be distributed among cells. However one still have to decide which set of nutrients can be inserted into the system. For a deeper characterization of given cancer populations, exo-metabolomic measurements to constrain the population boundary conditions would thus be needed. An exhaustive sensitivity analysis of scFBA results to boundary conditions is out of the scope of this work. However, it is interesting to compare the conditions in which the two major metabolites involved in cooperation (i.e., lactate and palmitate) are externally supplied to the population or must be produced endogenously.

Notably, we observed that uptake of exogenous palmitate does not affect the biomass production rate, indicating that no growth advantage is conferred by free availability of lipids. This result is in line with experimental evidence that cancer cells rely on *de novo* synthesis of palmitate-derived lipids [41]. However, in the baseline setting (no external palmitate supplied), we observed a group of cells that uptake the palmitate secreted by others (Fig. 5). We verified that, once internalized in those cells, palmitate is not processed by the beta-oxidation pathway, but directly contributes to the biomass synthesis, supporting the evidence reported in [10], that an exogenous source of fatty acids can substitute for *de novo* synthesis in promoting cell proliferation and attenuate the cancer-specific toxic effect of lipogenesis inhibitors. It has also recently been pointed out that a limited access to environmental lipids may render the cancer cells more sensitive to the inhibitors of lipogenesis [10]. In line with these findings, it can be observed, with regard to the LCPT45 population (Figure 6A), that a set of genes stops being lethal when exogenous palmitate is supplied. As expected, this set mainly includes genes directly involved in the synthesis of palmitate, namely: citrate synthase, fatty acid synthase and pyruvate dehydrogenase. The expansion of the plot in Figure 6A (left) shows that the latter (pyruvate dehydrogenase, ID: HGNC:8808) is lethal for each cell within the population. It should indeed be noted that when synthesis nor uptake of palmitate are possible, also cells that rely on the palmitate produced by other cells are necessarily affected.

**Figure 6:**
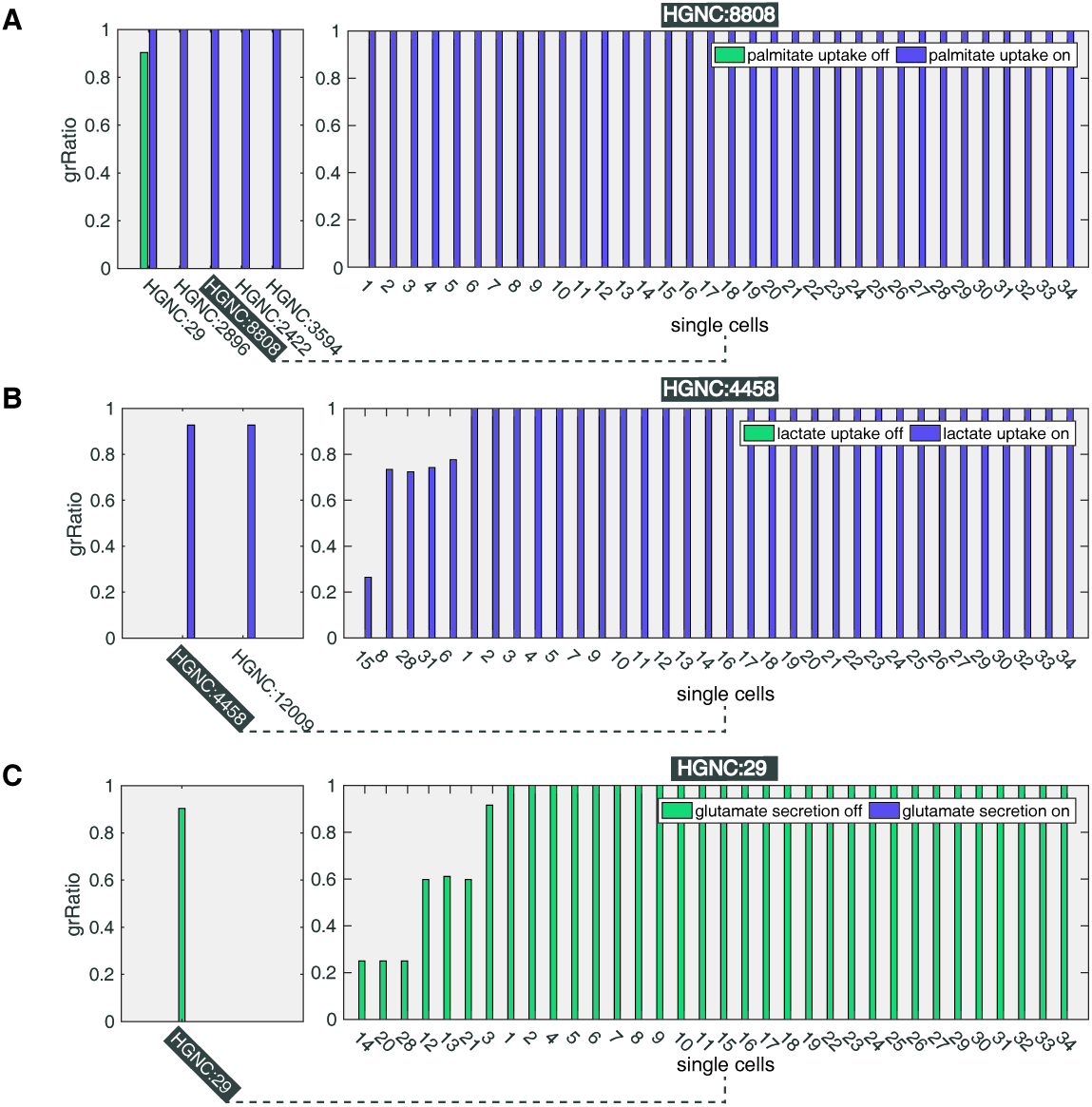
Impact of boundary conditions on gene-deletion predictions for LCPT45 dataset. A) Left: effect of gene deletions on the population growth rate, when exogenous palmitate uptake is allowed (purple bars) and when is not (green bars). Only genes with differential effect are reported. A missing bar indicate a lethal gene (*grRatio* = 0). Right: effect of the deletion of gene HGNC:8808 on the growth rates of each single-cell. B) Left: effect of gene deletions on the population growth rate when exogenous lactate uptake is allowed (purple) and when is not (green). Right: effect of the deletion of gene HGNC:4458 on each single-cell. C) Left: effect of gene deletions on the population growth rate when endogenous glutamate release is allowed (purple) and when is not (green). Right: effect of the deletion of gene HGNC:29 on each single-cell.

As opposed to palmitate, the metabolite lactate is not strictly required for growth. However, it can be observed in Figure 6B that the deletions of genes encoding for glucose-6-phosphate isomerase (HGNC: 4458) and for triosephosphate isomerase 1 (HGNC: 12009) - two important steps for the utilization of glucose through glycolysis - are not lethal when lactate uptake is allowed, suggesting that lactate may be able to replace glucose as carbon source. Interestingly, when lactate uptake is prevented, the plot expansion in Figure 6B (left) shows that the gene HGNC: 4458 is lethal in many but not all cells.

We remark that also the set of metabolites allowed to be released, e.g., in the blood affect the effect of gene deletions. Worth of note, if glutamate secretion is prevented, the deletion of the gene that encodes for palmitate secretion becomes lethal, as shown in Figure 6C. Remarkably, it has been reported that secretion of lipids facilitates tumour progression [28], whereas inhibitors of glutamate release have been proposed as new targets for breast cancer-induced bone-pain [14]. scFBA may enable to shed light on how the disposal of carbons through these two metabolites relates with the utilization pattern of exogenous nutrients.

## Discussion

To the best of our knowledge, we have presented the first attempt to solve the problem of reconstructing the single-cell fluxome, starting from single-cell transcriptomes, by taking into account environmental constraints, as well as cell-cell interactions. scFBA integrates single-cell transcriptomics data with (bulk) extracellular fluxes of the same cancer population, by means of a computational approach inspired to complex systems science [9]. Single-cell fluxes are computed as a function of transcriptome at the single-cell, while assuring that the constraints on mass-balance and on the rates of consumption/secretion of nutrients by the entire population are satisfied, along with the requirement for the tumour mass to grow. Although we do not explicitly model spatial organization, the constraints on single-cell transcriptome should implicitly preserve the information on the usage/secretion of most nutrients of each cell in its original position. Importantly, scFBA is able to point out the metabolic interactions that are established within a cell population.

As a proof of principle, we have successfully applied the methodology to LUAD and breast cancer datasets. We have shown how the integration of scRNA-seq greatly reduces the space of feasible solutions leading to metabolic growth of the overall population, which is prerequisite of tumour growth.

In the near future, single-cell transcriptome of thousands of cells will be obtained at the cost of a bulk experiment [39]. We have illustrated how, by translating sc-transcriptomes into sc-fluxomes, scFBA is valuable to extract features (i.e., the sc-flux values) from these data, in order to identify metabolic clusters of cells, which may be used to investigate other fingerprints of the cancer metabolic deregulation.

We have also shown how alternative objective functions, such as ATP maximization, may be investigated. sub-optimality may also be taken into account with sampling methods [8]. As shown in [7] the cost of popFBA computations and the space of feasible solutions to be explored linearly increases with the size of the template metabolic network and of the pool of sequenced cells.

Finally we have shown how constraints on the nutrient consumption and secretion rates (extracellular fluxes) of the specific sequenced population may affect scFBA predictions. As opposed to intracellular fluxes, (bulk) extracellular fluxes might be easily estimated, e.g., by approximation from metabolite concentrations in spent medium (exo-metabolome), when culturing patient-derived cells. Hence, datasets collected in the spirits of scFBA are needed to tune and fully prove the potential of scFBA to investigate intra-tumour heterogeneity starting from-omics data retrieved, e.g., from cancer biopses, paving the way to personalized population models of cancer metabolism.

## Acknowledgement

The institutional financial support to SYSBIO - within the Italian Roadmap for ESFRI Research Infrastructures - is gratefully acknowledged. LA received funding from FLAG-ERA grant ITFoC.

